# Enrichment of short mutant cell-free DNA fragments enhanced detection of pancreatic cancer

**DOI:** 10.1101/482745

**Authors:** Xiaoyu Liu, Lingxiao Liu, Yuan Ji, Changyu Li, Tao Wei, Xuerong Yang, Yuefang Zhang, Xuyu Cai, Ting Yang, Yangbin Gao, Weihong Xu, Shengxiang Rao, Dayong Jin, Wenhui Lou, Zilong Qiu, Xiaolin Wang

## Abstract

Analysis of cell-free DNA (cfDNA) is promising for broad applications in clinical settings, but with significant bias towards late-stage cancers. Although recent studies have discussed the diverse and degraded nature of cfDNA molecules, little is known about its impact on the practice of cfDNA analysis. Here we reported a new targeted sequencing by combining single-strand library preparation and target capture (SLHC-seq). By applying the new technology in plasma cfDNA from pancreatic cancer patients, we achieved higher efficiency in analysis of mutations than previously reported using other detection assays. SLHC-seq rescued short or damaged cfDNA fragments along to increase the sensitivity and accuracy of circulating-tumor DNA detection. Most importantly, we found that the small mutant fragments are prevalent in early-stage patients, which provides strong evidence for fragment size-based early detection of pancreatic cancer. Collectively, the new pipeline enhanced our understanding of cfDNA biology and provide new insights for liquid biopsy.

## MAIN TEXT

### Introduction

Pancreatic cancer is among the most lethal malignancies, with a 5-year survival rate of approximately 8%,(Siegel et al, 2017) and is predicted to be the second leading cause of cancer-related death by the year 2030.(Rahib et al, 2014) Early diagnosis and personalized treatment can greatly benefit patients with this disease.(Kamisawa et al, 2016; Lennon et al, 2014) Because targeted therapies have been reported to be highly effective for a subset of cancer patients with actionable mutations, characterizing pancreatic cancer at the genetic level holds great promise for precision medicine in the near future.(Garrido-Laguna & Hidalgo, 2015; Mardis, 2012; Witkiewicz et al, 2015)

However, genomic profiling of pancreatic cancer is challenging due to several limitations. First, pancreatic lesions often have a high stromal cell content, which may complicate the analysis of tissue sequencing.(Moffitt et al, 2015; Neesse et al, 2015; Waddell et al, 2015) Second, given the genetic instability and heterogeneity of pancreatic cancer, representative sampling is difficult for a comprehensive assessment of the genomic landscape of this disease.(Lee et al, 2016; Negrini et al, 2010; Notta et al, 2016) Multiple sampling using fine-needle aspiration (FNA) biopsies for pancreatic cancer is both unsafe and unfeasible in clinical practice.(Bournet et al, 2015; Fabbri et al, 2015) These challenges limit personalized medicine for pancreatic cancers. The possibility of detecting tumor-derived mutations in circulating cell-free DNA (cfDNA) provides a potential solution.(Bettegowda et al, 2014; Murtaza et al, 2013) cfDNA profiling can provide a more comprehensive snapshot of the genomic landscape of pancreatic cancers than tissue-based sequencing.(Aravanis et al, 2017; Zill et al, 2015) Moreover, the noninvasive nature of cfDNA analysis enables doctors to monitor disease progression.(Couraud et al, 2014; Murtaza et al, 2013; Zill et al, 2015) However, cfDNA analysis using large-scale next-generation sequencing (NGS) has been largely confined to advanced or metastatic cancers, in which the ctDNA (circulating-tumor DNA) fraction is higher and is more readily detectable.(Gorgannezhad et al, 2018; Haber & Velculescu, 2014) Detection of ctDNA in early-stage cancers is still filled with cautionary tales that highlight the technical challenges in this field. Recently, Velculescu *et al* reported improved targeted error correction sequencing (TEC-Seq) as an efficient approach for the detection of ctDNA in early-stage colorectal, ovarian, breast, and lung cancers, with a detection rate ranging from 59% to 71%.(Phallen et al, 2017) The massive parallel sequencing approach provides an applicable paradigm guiding cfDNA analysis in early-stage cancers.

However, the performance of existing approaches is limited in cfDNA analysis in early-stage pancreatic cancers. This limitation may be partially because of a lower ctDNA content in the circulation of pancreatic cancers.(Aravanis et al, 2017) Moreover, the biological features of ctDNA remain unclear, and this ambiguity may exacerbate the difficulty of detecting cancer-specific mutations in plasma of early-stage cancers. Recent advances have uncovered that cfDNA is a group of molecules with high diversification and degradation.(Burnham et al, 2016; Underhill et al, 2016) Conventional NGS or PCR approaches are not sensitive enough to capture the full diversity of these molecules, particularly degraded fragments with nicks in either strand. In 2012, Meyer *et al* introduced single-stranded library preparation for sequencing highly degraded DNA samples from an extinct archaic human.(Meyer et al, 2012) The single-strand approach has shown higher efficiency in the recovery of degraded and shorter DNA fragments.(Gansauge et al, 2017; Glocke & Meyer, 2017) Several recent studies have also shown that single-strand library preparation improved the recovery of cfDNA from cancer patients,(Snyder et al, 2016) transplantation recipients(Burnham et al, 2016) and maternal plasma.(Vong et al, 2017) It is of great significance to determine whether single-strand library preparation could indeed improve the performance of cfDNA analysis in clinical practice, especially in early-stage cancers.

Thus, we developed a single-strand library preparation and hybrid-capture-based cfDNA sequencing (SLHC-seq) approach and applied it to perform cfDNA profiling in a clinical cohort of 112 patients with pancreatic cancers and 28 healthy volunteers. SLHC-seq showed high sensitivity and specificity for detecting ctDNA in patients with pancreatic cancer across all stages. Using this new approach, the genomic landscape cfDNA was highly comparable to the results from direct tumor tissue sequencing. More importantly, single-strand library preparation efficiently enriched short cfDNA fragments (< 100 bp), which may prominently originate from degraded ctDNA fragments with nicks in either strand. We demonstrated that the ability to recovery these degraded fragments directly improved the sensitivity and accuracy of the detection of KRAS mutations in early-stage pancreatic cancer. Our observations highlight that the biologic features of ctDNA may ultimately determine the ability to detect precancerous or very small cancer lesions. The single-strand approach could be a powerful tool for large-scale mutation measurements of cfDNA and has the potential to be generalized to cfDNA analysis for other common cancers.

## Results

### Single-strand library preparation and hybrid-capture-based cfDNA sequencing

Based on the hypothesis that single-strand DNA library preparation is, in principle, more sensitive to recover the full spectrum of cfDNA fragments from plasma, we developed a single-strand library preparation and hybrid-capture-based cfDNA sequencing methodology (SLHC***-seq***, ***Fig. 1A,B)***. First, we selected the 62 most recurrently mutated genes in common cancer types according to the TCGA, ICGC database (International Cancer Genome Consortium et al, 2010) and large-scale cohorts relevant to pancreatic cancer sequencing.(Bailey et al, 2016; Moffitt et al, 2015; Waddell et al, 2015; Witkiewicz et al, 2015) Then, we designed hybrid-capture probes covering all exons of these 62 genes, which encompassed an approximately 211-kb region (***Table S1***). Theoretically, the single-strand library preparation should improve the conversion efficiency of degraded DNA fragments (especially fragments with nicks in either strand) compared with that of conventional methods. The following steps were similar to those in the targeted sequencing of cancer tissue. Bioinformatics analyses were performed to identify tumor-specific mutations and eliminate potential artifacts (Fig. ***S1)***. These analyses mainly included the following: (i) using unique molecular identifier to reduce possible contamination and systematic error; (ii) removing germline mutations according to the 1000 Genome database,(Clarke et al, 2012) and uncommon variant allele fraction (AF ≥ 40 %) were removed; and (iii) enrichment of likely cancer-specific mutations by removal of alternations not identified in common cancer types according to the COSMIC database(Forbes et al, 2011) (instances of overlapping mutations ≤ 2 were removed) (***Fig. S1***).

**Fig. 1.**
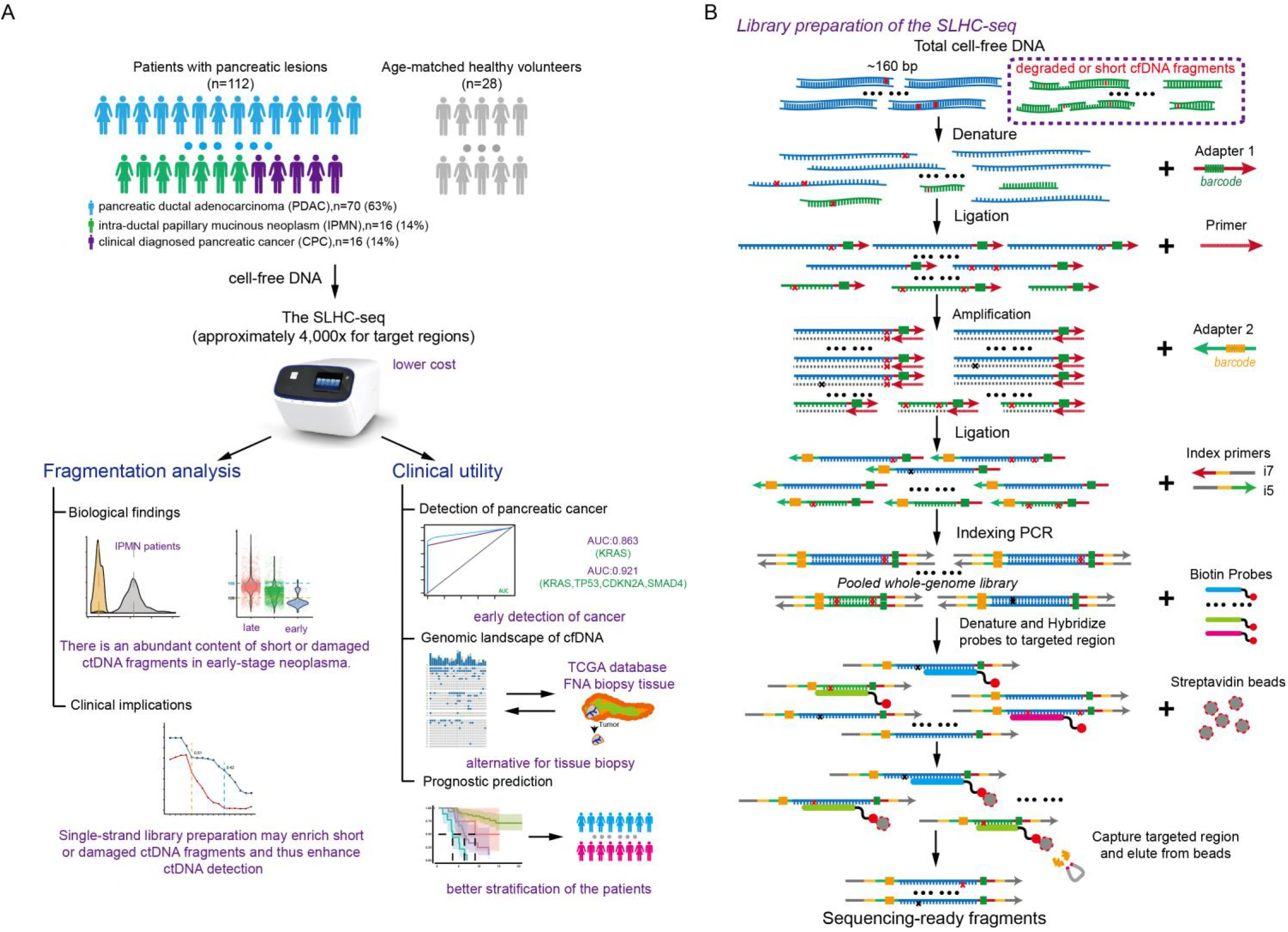
The study design and single-strand library preparation of cfDNA in conception. cfDNA fragments were first denatured into single-strand DNA (ssDNA) fragments. Then, the ssDNA fragments were ligated with a unique molecular identifier. The pre-library was enriched using hybridization and magnetic-beads capture and then subjected to high-throughput sequencing.

We next compared the performance of SLHC-seq with that of conventional NGS methods for detecting known cancer-specific mutations, including single nucleotide variants and small insertions and deletions, in reference genomic DNA. Generally, SLHC-seq and conventional approaches both showed high sensitivity and specificity in the analysis of mutations with higher fractions. Initially, with 1 ng of DNA input, SLHC-seq, compared with the conventional approach, showed a significantly higher efficiency for the detection of mutations in the reference genome (detection rate: 75% *vs.* 45%, respectively, *p<0.01, **Fig. S2***). Then, we evaluated the performance of SLHC-seq using a mixture of references at different dilutions. Initially, with 5 ng of DNA input, SLHC-seq identified over 50%, 70%, 92% and 92% mutations in 10, 5, 3,2-fold dilutions, respectively. Further analysis of the variant alleles achieved high concordance to the expected fraction of mutations (adjusted R^2^=0.94; *p<0.0001*, ***Fig. S2, Table S2***), as well as high sensitivity and specificity. Overall, the analytical sensitivity was 92%, 88% and 83% for detecting mutations using a threshold allele fraction of 0.5%, 0.2% and 0.1%, respectively. No false-positive mutations of the identified alleles were detected in the blank control using the SLHC-seq approach. According to the results, an allele fraction of 0.5% was determined as the threshold for identifying cancer-specific mutations in plasma cfDNA.

### Detection of KRAS mutations using SLHC-seq in the circulation of pancreatic cancers

Of the 112 included patients, 70 were histologically diagnosed as having pancreatic ductal adenocarcinoma (PDAC), 16 were intra-ductal papillary mucinous neoplasm (**IPMN**), and 16 were clinically diagnosed pancreatic cancer (**CPC**) ***(Fig. S3, Table S3)***. The median cfDNA concentration across the pancreatic patient group was 16.2 ng/ml (IQR: 9.3-25.9 ng/ml), which was significantly higher than that in the healthy individual group (median 3.2, *p<0.0001*, Wilcoxon-Mann-Whitney test). The total cfDNA concentration of the patients was independent of tumor stage and CA19-9 value, while it increased with radiological tumor sizes in patients with localized PDAC (spearman rho=0.4021, *p=0.0042*, ***Fig. S3***).

Detection of KRAS hotspot mutations in the circulation was often used for diagnosis of pancreatic cancer. We compared the performance of our approach in measuring KRAS mutations with that in previous reports using PCR- or NGS-based methods (***Table S4***). (Allenson et al, 2017; Berger et al, 2016; Brychta et al, 2016; Cheng et al, 2017; Cohen et al, 2017; Hadano et al, 2016; Kim et al, 2018; Kinugasa et al, 2015; Le Calvez-Kelm et al, 2016; Pietrasz et al, 2017; Sausen et al, 2015; Sefrioui et al, 2017; Takai et al, 2015) Across 13 high-quality studies, the detection rate of the KRAS mutations in plasma ranged from to 21% to 76% (***Fig. 2A***). In our cohort, which included a substantial proportion of early-stage disease patients, tumor-specific mutations were detected in 88% (99/112) of patients, with KRAS hotspot mutations detected in 70% [95% confidence interval (CI): 62-79%] of patients (***Fig. 2B***). Of note, SLHC-seq largely outperformed most reports using the ddPCR method in the detection of KRAS mutations in early-stage diseases. Using the SLHC-seq approach, the detection of KRAS mutations served as an efficient marker to distinguish PDAC from healthy individuals in our cohort [AUC (area under curve): 0.863, 95% CI: 0.830-0.898, ***Fig. 2C***]. By utilizing a combination of the KRAS, TP53, CDKN2A and SMAD4 genes, we achieved a higher performance in the diagnosis of PDAC (AUC: 0.921, 95% CI: 0.890-0.956, sensitivity: 80%, specificity: 100%, ***Fig. 2D***). Diagnostic accuracy was highest when using the full targeted panel, with an AUC of 0.951 (95% CI: 0.932-0.983) with an optimum specificity of 100% and sensitivity of 89%. Additionally, the detection of mutations using the full targeted panel also had the ability to distinguish IPMN from PDAC patients (AUC: 0.837, 95% CI: 0.821-0.979, specificity: 87.5%, sensitivity: 66.2%).

**Fig. 2.**
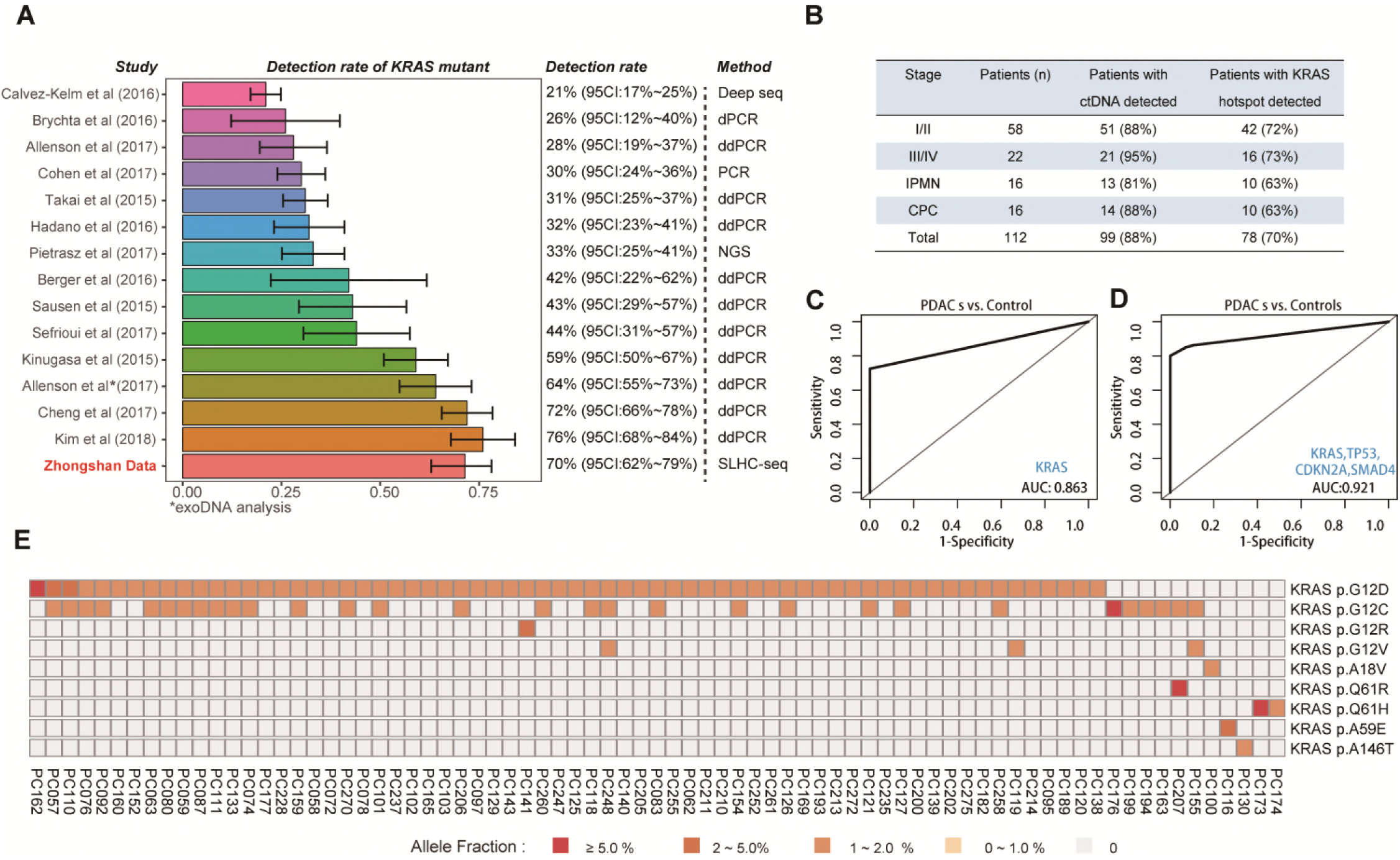
Detection of KRAS mutations in plasma of patients with pancreatic cancers. **(A)** Comprehensive analysis of cfDNA-based measurements of KRAS mutations in 13 previous reports. **(B)** ctDNA and KRAS hotspot mutations detected in cancer patients using SLHC-Seq. Diagnosis of PDAC using detection of mutations in **(C)** KRAS gene or **(D)** a combination of KRAS, CDKN2A, TP53 and SMAD4. **(E)** Heatmap is presented to illustrate the KRAS mutational heterogeneity in plasma ctDNA. Color scale of each square represents the allele frequency of mutations.

Of note, multiple KRAS hotspot mutations, including G12D (n= 67), G12C (n=30), were detected as well as other hotspot mutations at a lower prevalence (***Fig. 2E***). We identified 22 samples with a dominant KRAS clone concurrent with sub-clonal KRAS mutations with lower allele fraction, suggesting that these patients may harbor multiple KRAS mutations in tumor tissues.

### SLHC-seq enhanced detection of PDAC via enrichment of ultra-short fragments bearing KRAS hotspot alleles in early-stage disease

A pooled analysis of 96 patients and 28 healthy individuals show that the fragment length of total cfDNA between the two groups was similar (median: 161 *vs.* 162 bp). Utilizing single-strand library preparation, we explored the fragment length of cfDNA with a focus on the KRAS gene, given the extremely high prevalence of KRAS mutations in pancreatic cancers. Interestingly, a large fraction of patients with pancreatic tumors have mutated KRAS fragments below 100 bp, while the fragments containing wild-type KRAS alleles of these patients are homogenously approximately 160 bp in length (Fig. ***3A***). A pooled analysis of 78 patients showed that the ctDNA fragments bearing the KRAS mutant alleles, which originated from the pancreatic cancer, were significantly shorter than their wild-type counterparts (median: 135 *vs*. 164 bp, *p<0.0001*, ***Fig. 3B***).

**Fig. 3.**
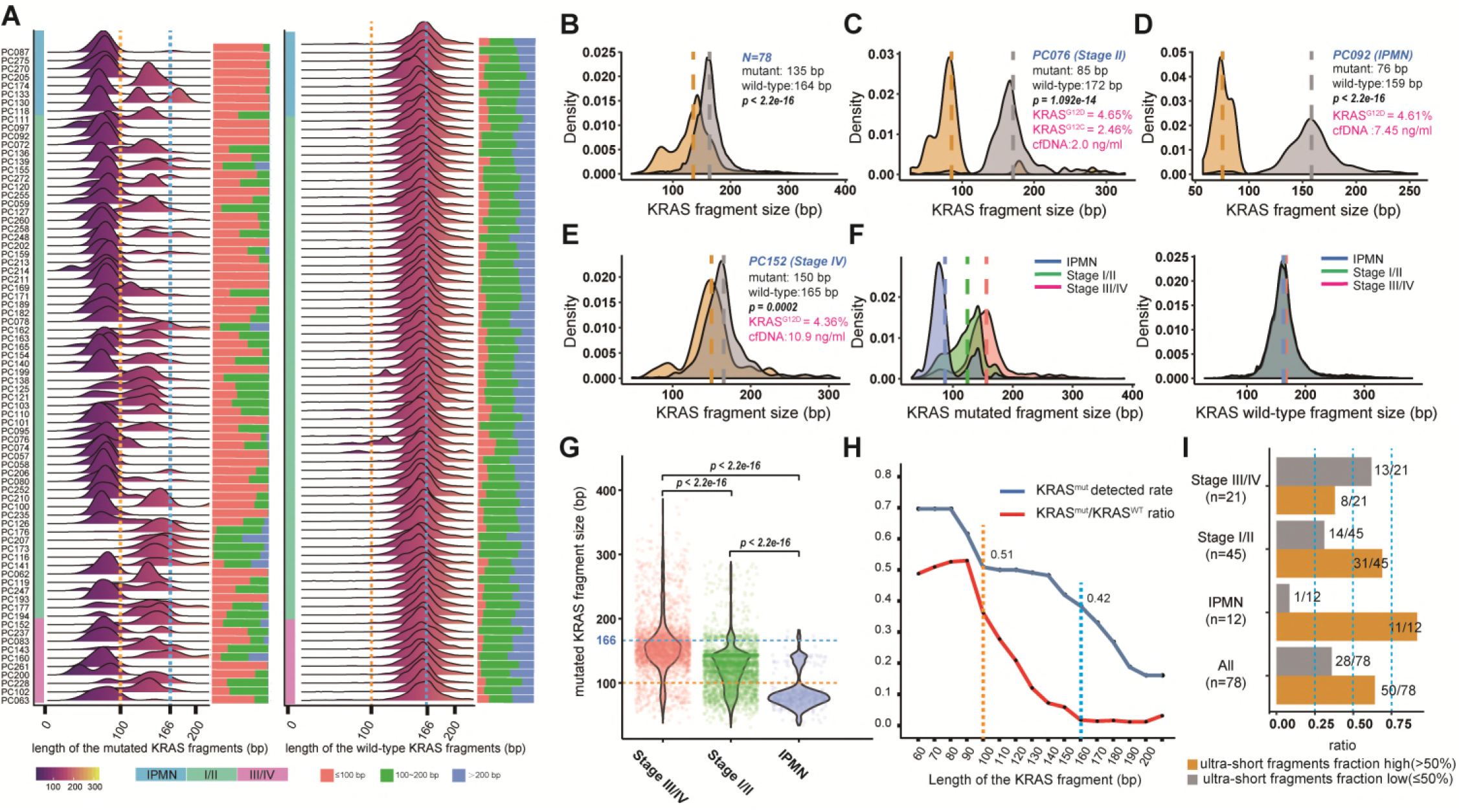
Ultra-short fragments containing the KRAS hotspot alleles in patients with pancreatic cancer. **(A)** Comprehensive view of fragment lengths bearing KRAS mutated and wild-type alleles derived from cell-free DNA deep-sequencing in patients with pancreatic cancer (n=78). (B) The fragment length of cfDNA bearing mutant KRAS alleles tended to be significantly shorter compared with DNA fragments bearing wild-type allele by densitometry in pancreatic cancer (n=78). The fragments bearing KRAS mutant alleles tended to be enriched in the region below 100 bp in patients (C) PC076 (stage II) and (D) PC092 (IPMN). (E) The distribution of KRAS mutated fragments included a large proportion of overlapping fragment sizes with wild-type KRAS fragments in advanced patients (PC152). (F) There was considerable discrepancy across IPMN (n=10), early-stage (n=42) and advanced (n=26) pancreatic cancer in terms of the length of KRAS mutated fragments by densitometry. (G) Violin plots representing the length of KRAS mutated fragments across different disease stages. (H) The ability of library preparation to enrich shorter fragments has a great impact on the detection of KRAS hotspot mutations in plasma (blue line), and the mutated-to-wild-type fraction of fragments bearing KRAS alleles reached the highest in the interval of 60-100 bp and declined sharply thereafter (red line). Ii) Ultra-short fragments (<100 bp) accounted for a quite high proportion of total KRAS mutated fragments in a substantial group of patients with pancreatic cancer.

Strikingly, the shortened fragment length was more pronounced in pre-cancerous IPMN and early-stage PDAC (stage I/II) patients. For example, in one case with IPMN (PC092), the fragments bearing the KRAS^G12D^ and KRAS^G12C^ allele were enriched in the region with a shorter length, and a dominant peak 75 bp in size (***Fig. 3C***). In another patient with stage II resectable PDAC (PC076), the distribution of fragments bearing the KRAS^G12D^ allele also showed significant deviation from the wild-type fragments with a dominant peak 85 bp in length (***Fig. 3D***). In contrast, in an advanced-stage patient (PC216) with a high mutated KRAS fraction (~25%), we observed a broad distribution of mutated fragments including a major proportion of fragments that overlapped with the wild-type fragments at a peak approximately 135-180 bp in length (***Fig. 3E***). The median length of the mutated fragment in IPMN (n=10), early-stage (n=58), and advanced-stage (n=10) PDAC was approximately 80, 140, 160 bp, respectively (*p<0.0001*, ***Fig. 3F-G***), while the wild-type counterparts had a length of ~160 bp across the subgroups (***Fig. 3F***). Of note, the relative wide distribution of wild-type fragments also included a small proportion of overlapping short fragments with the KRAS mutants, possibly as a result of the heterozygous features of KRAS mutations.

Subsequently, the length of fragments was binned into subgroups (60=0-60 bp, 70=71-70 bp, etc.), and the mutated-to-wild-type fragment ratio was calculated for each interval. The proportion of the fragments bearing KRAS mutations reached the maximum in the interval of 60-100 bp and sharply declined thereafter. The proportion of the KRAS mutated fractions declined to less than 5% for regions of 150-200 bp in length. Importantly, we found that the ability of library preparation methods to enrich these shorter mutated fragments had a significant impact on the performance of detecting KRAS mutations in plasma. The detection rate of KRAS mutations significantly decreased when setting the threshold at 100 bp or 160 bp (100 bp: 0.70 *vs*. 0.51, respectively, *p<0.05*; 160 bp: 0.70 *vs*. 0.42, respectively, *p<0.05*, ***Fig. 3H***). The ability to recover ultra-short fragments also affected the quantification of KRAS mutations in the plasma of patients with pancreatic cancer (Fig. ***3I***).

### Genomic landscape of cell-free DNA in patients with pancreatic cancer

We next explored the genomic landscape of cfDNA in pancreatic cancer using SLHC-seq. Generally, we identified 791 somatic mutations in the plasma of 88% (99/112) patients (approximately 4000-fold coverage, ***Fig. S4***). In contrast, 8 mutations were detected in the plasma of the healthy controls (n=28) without any known. oncogenic or truncating alleles. The average burden of cancer-related mutations was significantly higher in the plasma of patients with detectable ctDNA than that in healthy individuals (8.0 *vs.* 0.29 per individual, *p<0.01*).Mutations in the TP53, KRAS, CDKN2A and SMAD4 genes were detected in 86%, 79%, 70%, and 58% of plasma cfDNA samples, respectively. In addition to these driver genes, we also observed a high prevalence of mutated genes in the chromatin-remodeling pathway (KDM6A, AIRD1A, and PBRM1) and axon guidance (ROBO1, BOBO2) (***Fig. 4 A-B***). KRAS mutations (KRASG^12D^ and KRAS^G12V^) were the most recurrent mutations in the cohort, and we also observed R175H and R273H mutations of TP53, both frequently present across cancers. In addition, we observed that 13% (15/112) of patients harbored actionable mutations, including mutations of BRAF, ERBB2, PIK3CA, JAK2 and JAK3 genes.(Landrum et al, 2016) The BRAF^V600E^ mutation was detected in 3 (2.6%) patients with allele fraction ranging from 0.7% to 0.9%. JAK3^A573V^, recurrently mutated in hematopoietic malignancies, was observed in 2 (1.8%) patients. JAK2^V617F^ was detected in the plasma of a 67-year-old male patient independent of KRAS hotspots mutations (allele fraction: 9.1%, ***Fig. S4***). While several JAK1/JAK2 inhibitors have been reported to have efficacy in cancer patients with JAK2-activating mutations, these observations indicated that large-scale cfDNA tests may help to identify individuals who would benefit from genotype-driven therapy trials.(Hurwitz et al, 2015)

**Fig. 4.**
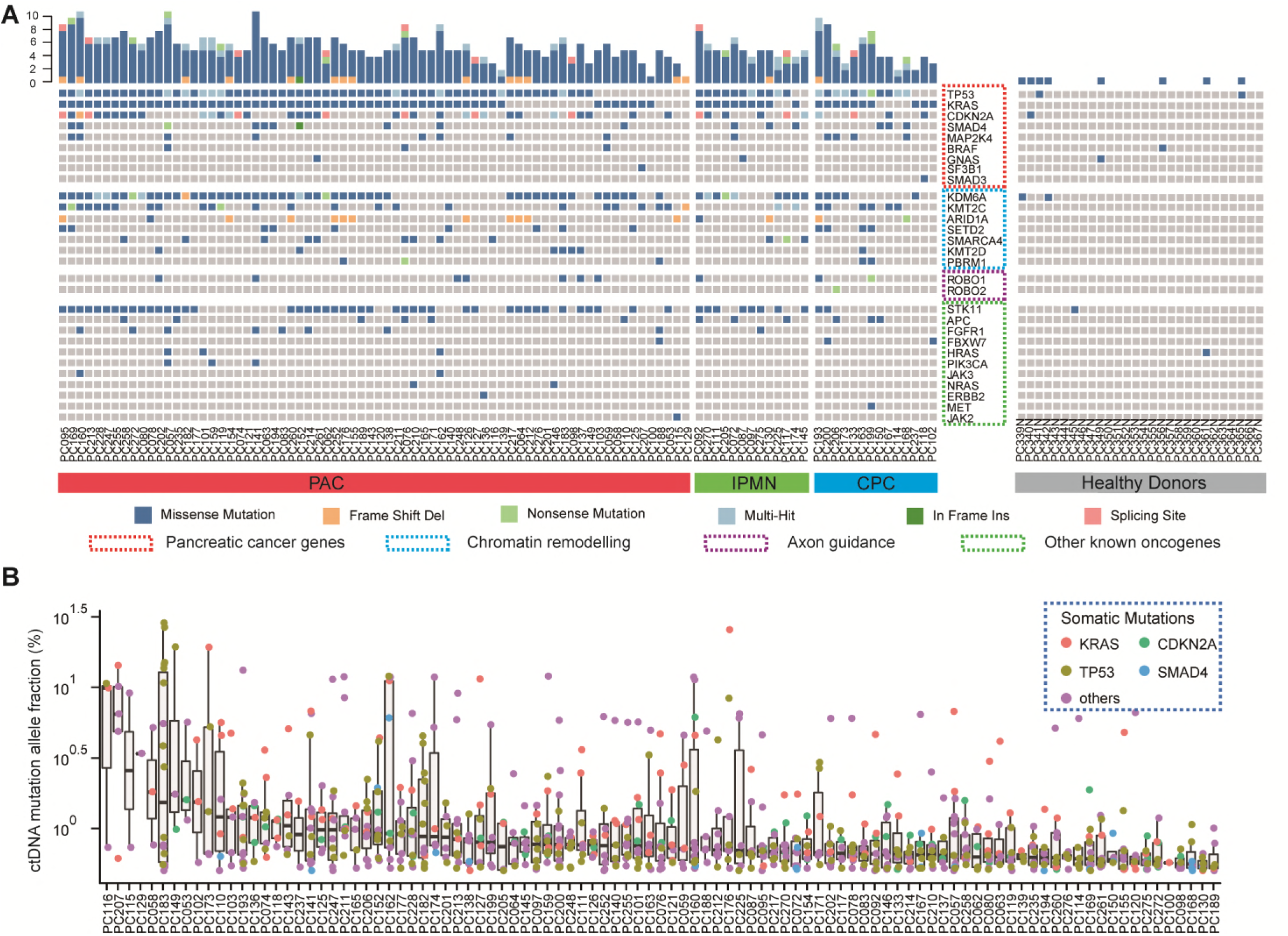
Genomic landscape of cfDNA in patients with pancreatic cancers. **(A)** Genomic landscape of cfDNA in patients with pancreatic cancers. A total of 792 somatic mutations were identified in the plasma of 88% patients. The mutations were highly prevalent in common pancreatic cancer genes and in genes involved in chromatin remolding, and axon guidance pathways. A total of 8 mutations were detected in the plasma of the healthy controls (n=28). The upper bars indicate the total mutation count for each sample. **(B)** The allele fraction of ctDNA mutation in each patients is shown; the box was sorted according to the median value of the mutational allele fraction in each individual. The colored dots represent mutations on different genes.

### Comparison of cfDNA profiling and tissue-based sequencing of pancreatic cancers

We next compared the cfDNA profiling results in our cohort and tissue-based sequencing data from 4 publicly available cohorts comprising 922 pancreatic cancer patients. For consistency, the top 15 mutated genes with prevalence over 5% in cfDNA were included in the final analysis. Generally, the cfDNA profiling in patients with detectable ctDNA was largely consistent with the results of the tissue-based sequencing database. The prevalence of KRAS, TP53, CDKN2A and SMAD4 mutations was highly correlated in plasma cfDNA and tissue-based cohorts (Fig. ***5A***). The mutational prevalence of the most commonly mutated genes in cfDNA was significantly associated with the mutational prevalence in tumor tissues (adjusted R^2^=0.456, *p=0.0057, **Fig. S5***). However, there was a significant discrepancy in the prevalence of KDM6A (67% *vs.* 13%*, p<0.001*) and STK11 (56% *vs.* 6%, *p<0.001*) between ctDNA profiling and tissue-based sequencing (***Figure 5A,B***). The prevalence of ROBO1 mutations was also not uniform across the tissue-based sequencing dataset. After the removal of KDM6A, STK11 and ROBO1, we observed a stronger correlation of the genomic landscape of cfDNA and tumor tissue profiling (adjusted R^2^=0.8723, *p=5.47e-06*, ***Fig. 5C***). The amino acid changes of KRAS predicted by cfDNA profiling were highly consistent with tissue-based sequencing (cBioPortal, ***Fig. 5D***).

**Fig. 5.**
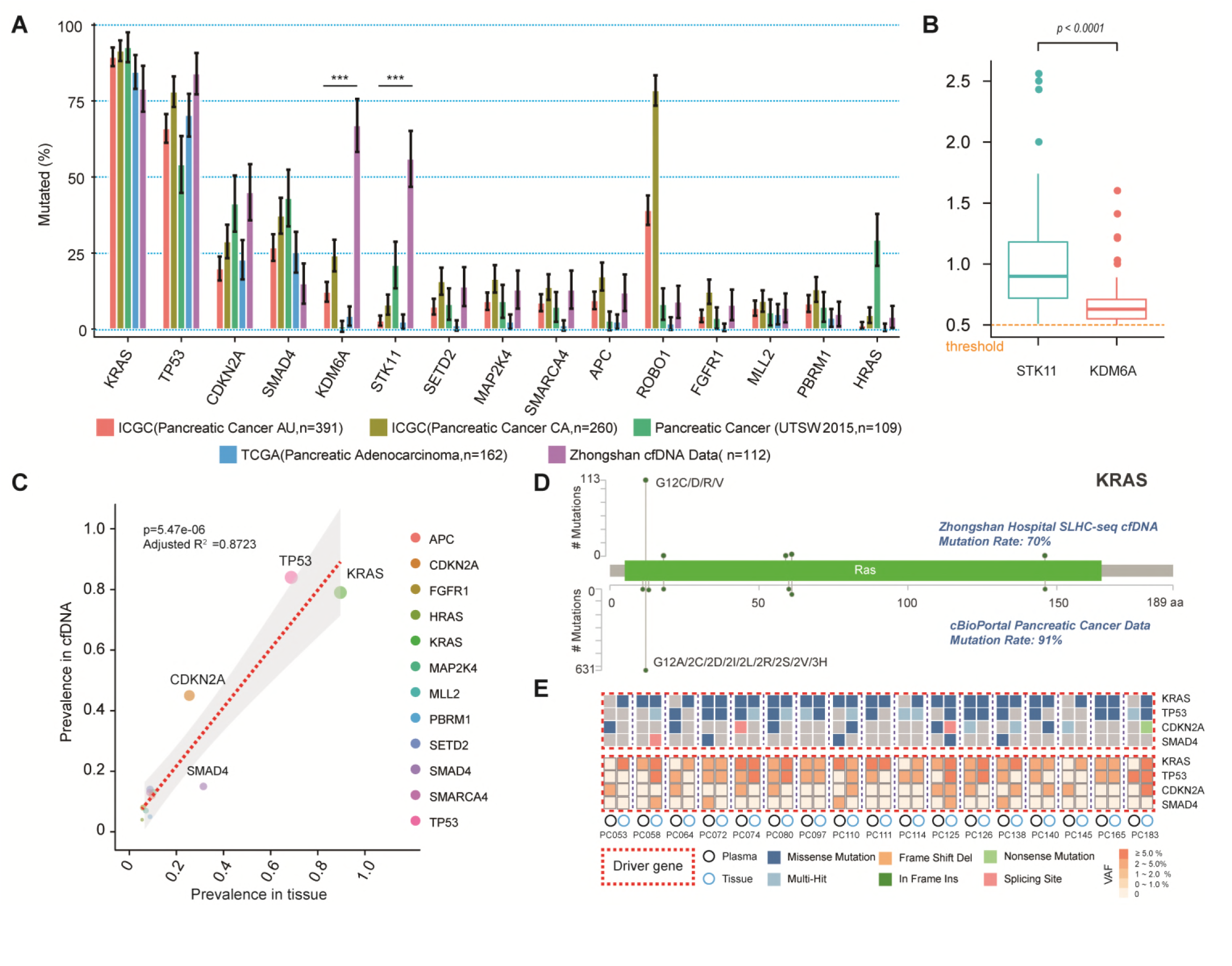
Genomic landscape of cfDNA or tumor tissue sequencing in pancreatic cancer. **(A)** Comparison of mutational prevalence in cfDNA and tissue-based cohorts (SNV/InDel only). The top 15 mutated genes in cfDNA are listed. **(B)** Comparison of the mutational allele fraction of STK11 and KDM6A in cfDNA. **(C)** Correlation between mutational prevalence in cfDNA versus tissue-based datasets (STK11, KDM6A and ROBO1 were removed). **(D)** The amino acid changes of KRAS predicted by cfDNA profiling versus tissue (cBioPortal). (E) Mutation profiling in cfDNA and paired biopsy tissue. The heatmap represents the mutation types, and the lower heatmap indicates summarized variant allele fractions (including clonal and subclonal) for each gene.

Additionally, we assessed the concordance of mutations in cfDNA and DNA from paired tumor tissues (n=17, ***Fig. S5***). Tumor cellularity ranged approximately from 1% to 15% in the biopsy tissue. The concordance of KRAS, TP53, and CDKN2A alternations between tissue DNA and cfDNA was 75.3%, 58.8%, and 41.2%, respectively (Fig. ***5E***, ***Table S2***). SMAD4 mutations were sporadically detected in 3 plasma samples and 2 tissue samples.

### ctDNA level and disease burden

In our study set, 74 patients were followed-up for a median time of approximately 2 years. The median overall survival of the patients was 15.9 months. Higher levels of CA19-9, CA125, CEA and total cfDNA concentration strongly correlated with poorer overall survival (***Table S5***).

Allele fraction of the mutations was previously reported as an indicator of plasma ctDNA burden. However, the concentration of total cfDNA is not specific to the specific cancer and is associated with conditions such as cancer-related inflammation. Thus, we estimated both the relative and absolute levels of KRAS mutations using the fraction of mutant-to-wild-type fragments and mutated ctDNA copies per ml in blood (Fig. ***6A***). The copies of KRAS mutated fragments (copies/ml) was associated with the mutant allele fraction and was also affected by total cfDNA quantity (R^2^=0.4456, *p<0.0001*, ***Fig. 6B***). A higher KRAS mutant fraction was associated with a poor prognosis (HR=2.1, 95% CI: 0.9-4.8, *p=0.077*, ***Fig. 6C***) but the difference did not reach significance. In contrast, the absolute quantification of KRAS mutations (copies/ml) directly marked patients with a worse prognosis to a good prognosis (HR=4.3, 95% CI: 2.1-9.1, *p<0.001*, ***Fig. 6D***). After adjusting for AJCC stage, the absolute quantification of KRAS mutations still directly marked patients with a worse prognosis (HR=2.9, 95% CI: 1.4-6.3, *p=0.007*, ***Fig. 6E-F***). In the multivariable analysis, the copy number was a significant marker of OS independent of AJCC Stage, CA19-9, CA125 and CEA level (HR: 3.3, 95% CI: 1.1-10.6; *p=0.037*, cox proportional hazards model, ***Table S6***). Together, our results suggest that accurate quantification of KRAS mutations in plasma may provide a useful marker for prognostic prediction in patients with pancreatic cancer across all stages.

**Fig. 6.**
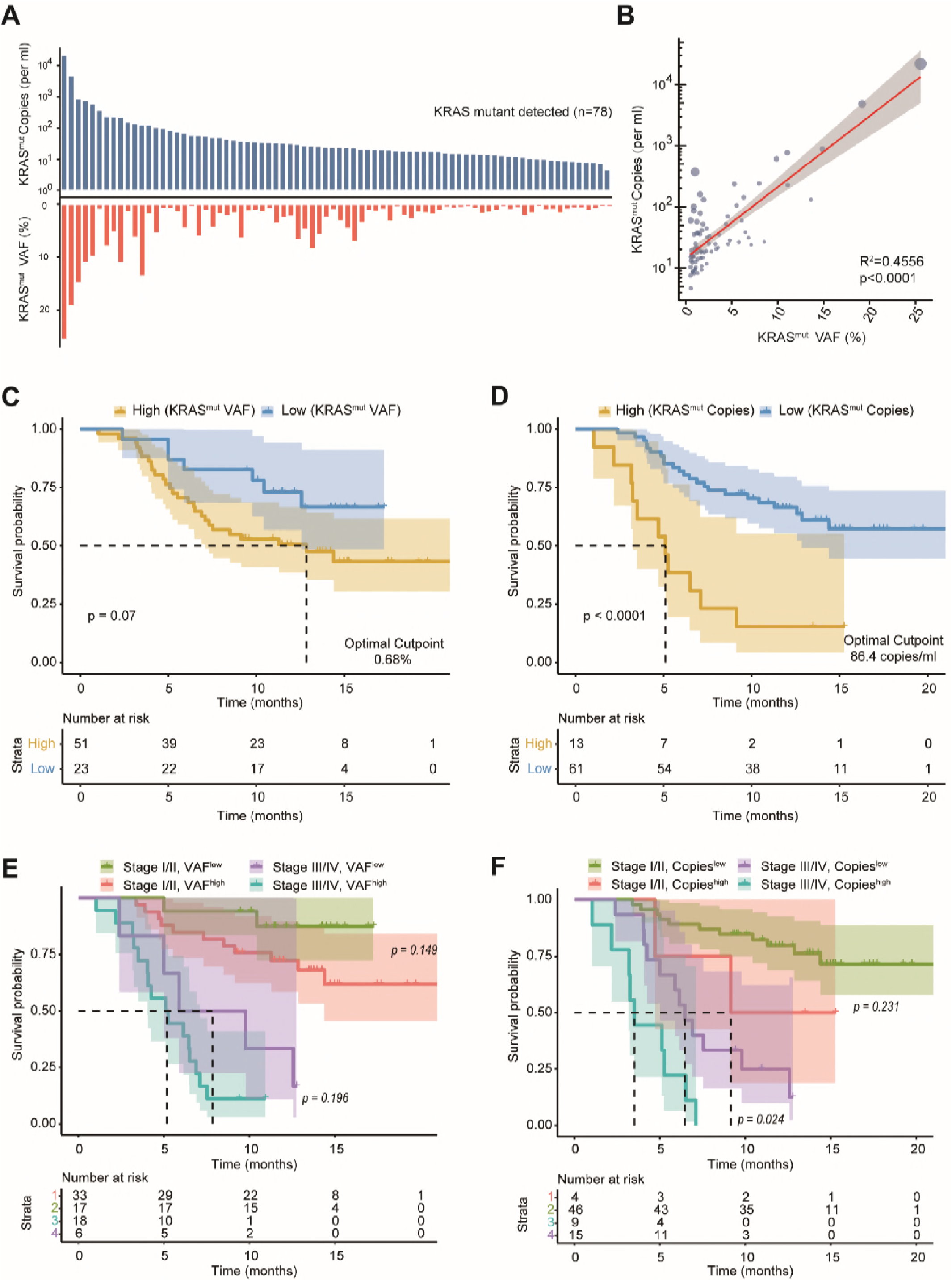
ctDNA level in plasma correlated with patient prognosis. **(A)** Relative and absolute levels of ctDNA bearing a KRAS mutation in each individual. **(B)** Quantitative correlation between the relative and absolute levels of KRAS mutations (the size of the blue dots represents the total cfDNA concentration in each case). The **(C)** relative and (D) absolute levels of KRAS mutation in plasma strongly correlated with the overall survival of the patients. **(E-F)** The absolute level of KRAS mutation in plasma was superior to the relative fraction for prognostic prediction together with AJCC tumor staging (edition 7).

## Discussion

Here by application of single-strand library preparation, we refined the cfDNA profiling in a large cohort of patients with pancreatic cancer across all stages. The cfDNA profiling of pancreatic cancer was highly comparable to tissue-based sequencing in terms of the mutational landscape. Based on previous research, we further revealed the different fragmentation patterns of ctDNA between early and advanced pancreatic neoplasms. Together with several current reports, the results demonstrate that the fragmentation pattern of ctDNA may largely affect the detection and quantification of ctDNA in the circulation. These findings facilitate a deeper understanding of ctDNA biology and support the validity and feasibility of single-strand library preparation for ctDNA analysis in early-stage cancers.

cfDNA analysis could provide a non-invasive approach for cancer diagnosis and surveillance of drug resistance and may also encompass the heterogeneity of tumor genomics.(Bettegowda et al, 2014; Phallen et al, 2017; Sausen et al, 2015; Zill et al, 2015) Recently, studies focusing on the Circulating Cell-free Genome Atlas (CCGA) have been performed to identify the genomic landscape of cfDNA in cancer patients and healthy participants.(Aravanis et al, 2017) This study will provide critically important knowledge regarding cfDNA profiling and improve the early diagnosis of cancers. Despite this promise, the identification of ctDNA signals is technically challenging, especially in non-metastatic solid tumors.(Aravanis et al, 2017) Conceptually, a benefit of our new approach is the adoption of a single-strand library preparation, which can recover the full spectrum of cfDNA fragments in plasma and thus improve the applicability for the detection of tumor-specific mutations in early-stage cancers. Here, we achieved an effective sensitivity in ctDNA detection across all stages of pancreatic cancer with only approximately 4,000-fold sequencing coverage. Of note, the detection of KRAS mutations in the plasma of pancreatic cancer patients has been studied for a long time, but with significant bias towards late-stage patients. Using this new approach, KRAS hotspot mutations were detected in over 70% of patients in our cohort, which comprised 66% of patients with precancerous or early-stage disease, while no hotspot mutations were detected in the circulation of healthy individuals. The performance of SLHC-seq in identifying ctDNA was highly comparable to that of ddPCR-based methods or NGS with ultra-deep sequencing depth (10,000-fold coverage or more). The high sensitivity and reliability of SLHC-seq make it highly suitable for early cancer diagnosis and intervention, and its ability for large-scale parallel sequencing of multiple and broad genomic regions also enables the quantitative surveillance of tumor-derived mutations that may evolve during early tumorigenesis, tumor progression, and drug resistance.

Next, we tried to analyze the fragmentation pattern of cfDNA molecules to find out why SLHC-seq may improve the detection of tumor-derived signals. In previous reports, the periodical shortening pattern of ctDNA fragments has been well described in xenograft animal models or in patients with late-stage cancers.(Jiang et al, 2015; Underhill et al, 2016). In two important studies published recently, researchers found that size-based analysis of cfDNA fragments improved the detection of tumor-derived DNA fragments (Mouliere et al, 2018a; Mouliere et al, 2018b). Compared with their experimental pipelines, the advantages of the single-strand library preparation were to recover the full spectrum of cfDNA fragments. First, the different shortening pattern of cfDNA and ctDNA in our results was consistent with these previous reports. Then, as the emergence of KRAS hotspot mutations has been globally recognized as the initialization of pancreatic neoplasia, analyzing fragments bearing KRAS oncogenic mutations provides an ideal model for further exploring the biological features of ctDNA during early tumorigenesis. Here, we identified a substantial subset of KRAS mutated fragments with a length less than 100 bp, and this phenomenon was more pronounced in early-stage disease. Strikingly, almost all fragments bearing KRAS mutations were enriched at the region of less than 100 bp in length in quite a large portion of patients. The enrichment of ultra-short fragments (<100 bp) in ctDNA has hardly been reported before. Based on previous reports, we thought these observations could be more reasonably explained using a ctDNA model with widespread DNA damage (particularly with nicks in either strand), rather than a theory of efficient enrichment of ultra-short fragments.(Burnham et al, 2016; Gansauge et al, 2017; Snyder et al, 2016) While conventional library preparation worked poorly in the recovery of damaged fragments, the single-strand approach retains more biological information from those damaged ctDNA fragments.(Gansauge et al, 2017; Meyer et al, 2012; Snyder et al, 2016) The ability to enrich these damaged ctDNA fragments, which may recover quite a high proportion of ctDNA in early-stage cancers, directly affects the detection and quantification of ctDNA in circulation. As ctDNA is presumably shed from tumors either through necrosis or apoptosis, we propose a plausible hypothesis to explain the different fragmentation pattern of ctDNA between early- and late-stage cancers. ctDNA in early-stage cancers mainly originates from cancer cells that have disintegrated via apoptosis and immune clearance procedures, in which the endonuclease activation and uptake of macrophages may account for the widespread damage in DNA fragments. In advanced-stage cancer, hypoxia-related necrosis occurs due to increasing tumor size and deficient vascularization. Large amounts of tumor-derived DNA fragments may not be efficiently degraded through endogenous and exogenous pathways. These less degraded DNA fragments shedding into the circulation could be detected via conventional approaches while the highly degraded fragments could not. This hypothesis may, in principle, provide a plausible explanation for the low detection rate of KRAS mutations in early-stage pancreatic cancer even using extremely sensitive approaches. Collectively, our findings provide strong evidence that the biologic features of ctDNA may ultimately determine the ability to detect precancerous or very small cancer lesions.(Phallen et al, 2017)

Another import aspect of our study was to explore the utility of SLHC-seq in clinical practice. First, SLHC-seq improved the detection of KRAS hotspot mutations in early-stage pancreatic cancers. Then, by utilizing SLHC-seq, the genomic profiling of cfDNA showed high similarity to the genomic alternations identified in 4 tissue-based sequencing databases of pancreatic cancer. In a subset analysis of 17 paired plasma and biopsy tissue samples, cfDNA profiling accurately reproduced the landscape of driver genes (KRAS, TP53 and CDKN2A) highly mutated in biopsy tissues. As biopsy tissues of pancreatic lesions are not routinely accessible, the high concordance of the genomic landscape between cfDNA and tumor tissues suggested that highly efficient cfDNA analysis could complement tissue-based sequencing in clinical practice. Further, cfDNA analysis theoretically provides more comprehensive information on tumor-derived mutations that may otherwise be missed due to the selection bias of tissue sampling and thus provides new insights into cancer biology. In our cohort, we observed mutational heterogeneity, as shown by the presence of distinct KRAS hotspot mutations within a cfDNA sample. Though tumor heterogeneity has always been emphasized in oncology, sub-clonal mutations may be missed due to technical limitations and the selection bias of current approaches in tissue analysis. Through the application of multiple sampling and microdissection, recent studies have identified polyclonal KRAS mutations in the tissue of papillary mucinous neoplasms(Tan et al, 2015; Wu et al, 2011) and ductal adenocarcinoma of the pancreas(Cancer Genome Atlas Research Network. Electronic address & Cancer Genome Atlas Research, 2017). Complementing this progress, our observations strongly indicated that polyclonal origin may be a common feature during the formation of pancreatic tumors, as polyclonal KRAS mutations in approximately 20% patients with pancreatic neoplasm have been observed. As previous researchers have identified that distinct KRAS mutations play different roles in the oncogenic pathway of lung cancer, further studies are needed to uncover the role of multiple RAS pathways in pancreatic cancer. At last, ctDNA fragments bearing KRAS hotspot alleles are directly released from the tumor. We demonstrated the accurate quantification of KRAS mutations in plasma may provide a useful marker for prognostic prediction in patients with pancreatic cancer across all stages.

Several limitations of this study should be emphasized. First, alterations originating from a hematopoietic clone may confound the tumor-specific mutations, which may lead to differences between the cfDNA profiling and tissue-based sequencing and over diagnosis. Going forward, investigations are needed to determine whether these mutations are clinically meaningful over time in healthy individuals. Second, though we identified a subgroup of patients with potentially actionable mutations, correlation between clinical drug response and detection of targeted mutations still need further investigation. Third, experimental and analytical approaches are still needed to avoid the cross contamination of probes with of homologous of target regions. Lastly, sensitivity of ctDNA detection may be further improved by sequencing with deeper depth or repeated sampling.

## Conclusions

Profiling somatic alternations has become routine for cancer patients, but the non-invasive nature of cfDNA analysis makes it promising for broad application in terms of clinical use. We introduced a unique application of single-strand library preparation to recover degraded or short cfDNA fragments in conception. Here we refined the genomic landscape of cell-free DNA in patients with pancreatic cancer, and found strong evidence that fragmentation pattern of ctDNA may profoundly affect the detection of precancerous or very small cancer lesions. We further demonstrated the clinical significance of this approach that high efficient cfDNA analysis could be an alternative to tissue-based sequencing and efficient biomarkers for pancreatic cancer in clinic. Together with other recent advances, our results provide a deep understanding of cfDNA biology and may promote the development of new adaptive approaches for ctDNA-based testing in the future.

## Materials and Methods

### Experimental Design

The design schematic is presented in ***Fig. S1***. A total of 112 patients with pancreatic lesions were recruited at Zhongshan Hospital Fudan University from July 2015 to May 2016. In addition, 32 age-matched healthy individuals were recruited as the control group. A total of 10 ml of treatment-naive blood from each individual was collected using K_2_-EDTA tubes (BD, Cat# 367863) and processed within 4 hours. Formalin-fixed, paraffin-embedded (FFPE) tumor tissues (4 samples from FNA biopsy and 13 from surgical resection) were collected from the Department of Pathology at our center, and a single sample of resected tissue (n=13) was collected from each patient without further dissection, representing the ‘mimic’ for biopsies. Patients were followed up every 3 to 5 months by a full-time clinical investigator as previously described.(Liu et al, 2016; Zhou et al, 2017)

#### Ethics approval and consent to participate

The study protocol was approved by the Ethics Committee of Zhongshan Hospital Fudan University (Approval No. B2014-098 and B2017-048). This included consent to publish from the participant to report individual patient data in our report.

#### Single-strand library preparation, targeted sequencing and bioinformatics analysis

The full details of the DNA extraction, library preparation, NGS and bioinformatics metrics are presented as below. Generally, cfDNA was first denatured into single-strand fragments after end repair. Then, these fragments were processed for library construction and hybrid-capture of targeted regions covering all exons of these 62 genes, which encompassed approximately a 221-kb region (***Table S1***). We achieved approximately 4,000-fold coverage of sequencing. **FreeBayes** software (version 1.1.0-3) was used to call SNV/INDEL variants. The datasets of tissue-based sequencing were obtained from the to The Cancer Genome Atlas (TCGA, cBioPortal dataset) and International Cancer Genome Consortium (ICGC, Supplementary Table).(Cerami et al, 2012; Gao et al, 2013) R packages (‘maftools’ version 3.5.0)(Mayakonda & Koeffler, 2016) and custom scripts were used for further annotation, filtration and visualization of the dataset.

#### Extraction and preparation of cfDNA and genomic DNA

The clinical cohort and tissue bank was previous registered (http://www.chictr.org.cn/, ChiCTR-ROC-14005393). cfDNA was isolated using the QIAamp Circulating Nucleic Acid Kit (QIAGEN, Cat#55114) from 5~10 ml of plasma samples. Concentration of the total cfDNA was measured using the Qubit™ dsDNA HS Assay Kit (Thermo Fisher Scientific, Cat# Q32854), and the quality was analyzed using the Agilent High Sensitivity DNA Kit (Cat# 5067-4626) on an Agilent 2100 Bioanalyzer (Agilent Technologies). cfDNA samples with excessive high molecular weight DNA content were excluded from further processing. Genomic DNA of tissue samples was isolated using the Qiagen QIAamp DNA FFPE Tissue Kit (Qiagen, Cat# 56404) following the manufacturer’s instructions. The concentration of the genomic DNA was measured using a Nanodrop 2000 (Thermo Scientific). Genomic DNA was fragmented to an average size of 200 bp using M220 Focused-ultrasonicator™ (Covaris, Inc.) following the manufacturer’s instructions.

##### Pre-library construction

AnchorViola^TM^ pre-library construction was performed using AnchorDx SmartVisio^TM^ Library Prep Kit (AnchorDx, Cat# A0UX00671) and AnchorDx SmartVisio^TM^ Indexing PCR Kit (AnchorDx, Cat# A2DX00687). End repair of fragmented gDNA or intact cfDNA was performed using the VEE1 enzyme at 37°C for 30 minutes. The DNA was then denatured at 95°C for 5 minutes and snap chilled on ice. The VLE1 and VLE2 enzymes were used at 37°C for 30 minutes. First, amplification was immediately performed to generate reverse complementary DNA molecules using the VAB1 PCR master mix with the following PCR program: 1 cycle of 98°C for 30 s; 4 cycles of 98°C for 15 s, 60°C for 30 s, and 72°C for 30 s; and 1 cycle of 72°C for 5 minutes. Amplified DNA was purified using the AMB1 Magnetic Beads and eluted in 20 μl of EB buffer. Another round of 3’ end adaptor ligation was performed using the VSE1 and VSE2 enzymes at 37°C for 30 minutes, followed by indexing PCR (i5 and i7) using the VIB1 PCR master mix with the following PCR program: 1 cycle of 95°C for 3 min; 7 to 11 cycles (cycle number depending on the input DNA mass) of 98°C for 15 s, 60°C for 30 s, and 72°C for 30 s; and 1 cycle of 72°C for 5 minutes. The indexing PCR products were purified using IPB1 Magnetic Beads. The purified PCR products were whole genome libraries (pre-libraries), and their concentration was measured using the Qubit™ dsDNA HS Assay Kit. Pre-libraries less than 500 ng were disqualified for subsequent target enrichment.

##### Target enrichment

Target enrichment was performed using the AnchorDx SmartVisio^TM^ Target Enrichment Kit (AnchorDx, Cat# A0UX00693). A total of 1000 ng of pre-libraries from up to 4 samples was pooled for one target enrichment using our custom-made panel focusing on 62 genes. Briefly, HE, HBA and HBB blocking reagents were added to the 1000 ng pooled pre-libraries and completely dried using a heated spin vacuum. The mixture was reconstituted in 7.5 μl of VHB1 hybridization buffer plus 3 µl of VHE1 hybridization enhancer. The mixture was next denatured at 95°C for 10 minutes and immediately transferred to a hybridization oven preheated to 47°C. The capture probes were added, and the mixture was quickly mixed and transferred to a thermocycler for hybridization incubation following the manufacturer’s protocol. After hybridization, target DNA libraries captured by the biotinylated probes were pulled down using M270 Dynabead streptavidin beads (Thermo Fisher Scientific, Cat# 65306). Briefly, 30 μl of M270 beads were washed twice with 1× Binding Wash Buffer, and the supernatant was removed. Pre-library pools were added and mixed thoroughly with the beads by repeated pipetting, and the mixture was incubated on a rotator at 47°C for 45 minutes. After incubation for the probes to bind to the beads, 100 μl of 1× Transfer Buffer pre-warmed to 47°C was added to the mixture, and the whole content was transferred to a magnetic stand. The supernatant was quickly removed and discarded as soon as it turned clear, and the beads were washed twice using 1× Stringent Wash Buffer pre-warmed to 47°C. The beads were then washed once with 200 μl of room temperature 1× Wash Buffer I, once with 1× Wash Buffer II and then once with 1× Wash Buffer III. The enriched libraries bound on the beads were further amplified with P5 and P7 primers using the KAPA HiFi HotStart Ready Mix (KAPA Biosystems, Cat# KK2602) with the following PCR program: 1 cycle of 98 °C for 45 s; 12 cycles of 98°C for 15 s, 60°C for 30 s, and 72°C for 30 s; and 1 cycle of 72°C for 1 minute. PCR products were purified with Agencourt AMPure XP Magnetic Beads (Beckman Coulter, Cat# A63882) and eluted in 40 μl of EB buffer. The concentration of the enriched library was measured using the Qubit dsDNA HS Assay.

##### Read mapping and SNV/INDEL analysis

Sequencing adapters and 3’-low quality bases were trimmed from raw sequencing reads using Trim Galore, version 0.4.1 (https://github.com/FelixKrueger/TrimGalore). After adapter trimming and quality control, reads were mapped to the UCSC hg19 reference genome using BWA MEM (version 0.7.13) using the default settings. Next, custom scripts were applied to remove PCR duplications based on information regarding unique molecule identifiers (UMI). Finally, **FreeBayes** (version 1.1.0-3) was applied to call SNV/INDEL variants using the settings “-F 0.001–standard-filters–haplotype-length 0–min-coverage 100-C 5–report-all-haplotype-alleles”. Of note, the BAM was further checked using Integrative Genomics Viewer (IGV) to remove potential artifacts in TP53 and EGFR genes. The R packages ‘maftools’ (version 3.4) ‘ggplot2’(version 2.2.1) and custom scripts were used for further analysis and visualization of the data (R version 3.4.0).

##### Identification the fragment length of cfDNA fragments bearing KRAS mutated or wild-type alleles

First, reads were cleaned by trimming adapters and low-quality bases. Next, cleaned reads were mapped to human reference genome (hg19) using BWA MEM, followed by unique molecule index (UMI)-based deduplication using our custom script. Finally, somatic variants were identified using **Freebayes** with the following parameters [-F 0.001 –standard-filters –min-coverage 100 –report-all-haplotype-alleles -C 5; depth >=500, frequency >=0.5%,AO>=5(for cfDNA analysis); depth >=500,frequency >=2%,AO>=5 (for tissue analysis)]. For a variant site of interest, reads were grouped into REF and ALT by the genotype at the site, and insert size information was computed from the size field of the bam file after deduplication.The estimation of number of copies of ctDNA bearing KRAS mutated alleles was mainly based on the assumption that there are 3.3 pg DNA per haploid copy of genome, the formula was used for calculating mutant copies per ul of plasma through NGS method:

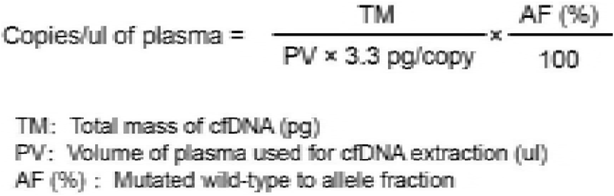

#### Statistical analyses

Chi-square test was used to analyze differences in categorical variables, while the Wilcoxon-Mann-Whitney test was used for continuous variables. Receiver operating characteristics (ROC) were used to evaluate the diagnostic power. In the univariate survival analysis, maximally selected rank statistics were used to determine optimal cutoffs for continuous variables, and the Kaplan-Meier method was next used to draft survival curves (log-rank test). Cox proportional hazards model was used to calculate the hazard ratio (HR) in univariate or multivariable survival analysis. Two-sized p-values less than 0.05 were considered significant. All statistical analyses were performed using R statistical project (version 3.4.0).

## Acknowledgments

We thank Dr. Yang Zhou, Dr. Junhao Li, Dr. Zhuqin Cao and Dr. Hao Liu for sample collection and follow-up of the patients. We thank the Author services from Springer Nature for English language editing.

## Funding

This work is supported by NSFC Grants (#91432111, #31625013, Recipient: ZLQ), Rong-Chang Charity Fund for Pancreatic Cancer Research (Recipient: XLW), Zhongshan Hospital Fund for Young Scholars (#2015ZSQN31, Recipient: CYL), National High Technology Research and Development Program of China (No.S2015AA020405, Recipient: WHL).

## Author contributions

Study concept and design: XYL, XLW, ZLQ, WHL; Acquisition of data: XYL, LXL, XRY, GYB, DYJ, SXR, YFZ, WHL; Analysis and interpretation of data: XYL, WHX, Drafting of the manuscript: XYL; Critical revision of the manuscript for important intellectual content: ZLQ, YBG, XYC, WT; Statistical and bioinformatic metrics: XYL, WHX; Technical, or material support: WHL, CYL, YFZ, YBG, CYL, LXL, TY; Study supervision: ZLQ, XLW, WHL.

## Conflict of interest

The authors declare that they have no conflict of interest.

## Data and materials availability

Mutation calls from FreeBayes and annotation are listed in Table S2. All other relevant data are within the paper and its supplementary files. According to ethical statements, interested researchers may contact Dr. Zilong Qiu to obtain raw human sequencing data.

## Supplementary Figure

**Fig. S1. Schematic of the study design and bioinformatics metrics. (A)** A general working flow of the study. **(B)** The bioinformatic schematic of the study.

**Fig. S2. The performance of SLHC-seq and conventional NGS approach for analysis known cancer-specific mutations in reference. (A)** The known cancer-specific mutations in reference genome [HD734, Tur-Q7 (1.3% Tie) standard DNA]. **(B)** Detection of somatic mutations using SLHC-seq and conventional NGS approach with different HD734 input. **(C)** Detection of somatic mutations using SLHC-seq with different HD734 dilutions. Analysis concordance of the variant alleles fraction (VAF) identified using SLHC-seq, conventional NGS approach **(D-E)** and the expected fraction of mutations.

**Fig. S3. Total cfDNA concentration in patients with pancreatic lesions and healthy controls.** 112 patients **(A)** and 32 healthy controls was included in the cohort. Total plasma cfDNA concentration in **(B)** patients with pancreatic lesions and healthy control group, **(C)** IPMN and PAC subgroups and **(D)** across cancer patients with different tumor burden. The distribution of cfDNA concentration of each individual is presented as hollow dots. **(E)** Total cfDNA concentration was correlated with radiological tumor size in patients with localized pancreatic malignancies (n=49).

**Fig. S4. The sequencing depth and potential actionable mutations.** The mean bait coverage **(A, B)** and the mean target coverage **(C, D)** were evaluated using the Hsmetrics function in the Picard toolbox version 2.5.0, and plotted in the descending order for each sample group, with the median and IQR shown as legends. **(E)** Potential actionable mutations detected in plasma cfDNA in patients with pancreatic cancers.

**Fig. S5. Comparison the genomic landscape of cfDNA and tissue-based sequencing. (A)** The mutational prevalence of the most commonly mutated genes in cfDNA was significantly associated with the mutational prevalence in tumor tissues. **(B)** Schematic illustrating comparison genomic landscape of cfDNA and paired tissue using SLHC-seq. **(C)** CT-guided fine-needle biopsy for patients with pancreatic lesion.

## Supplemental Tables

**Large excel datasets of Table S1, Table S2 and Table S3 were uploaded as independent files in the submission system.**

**Table S4.**
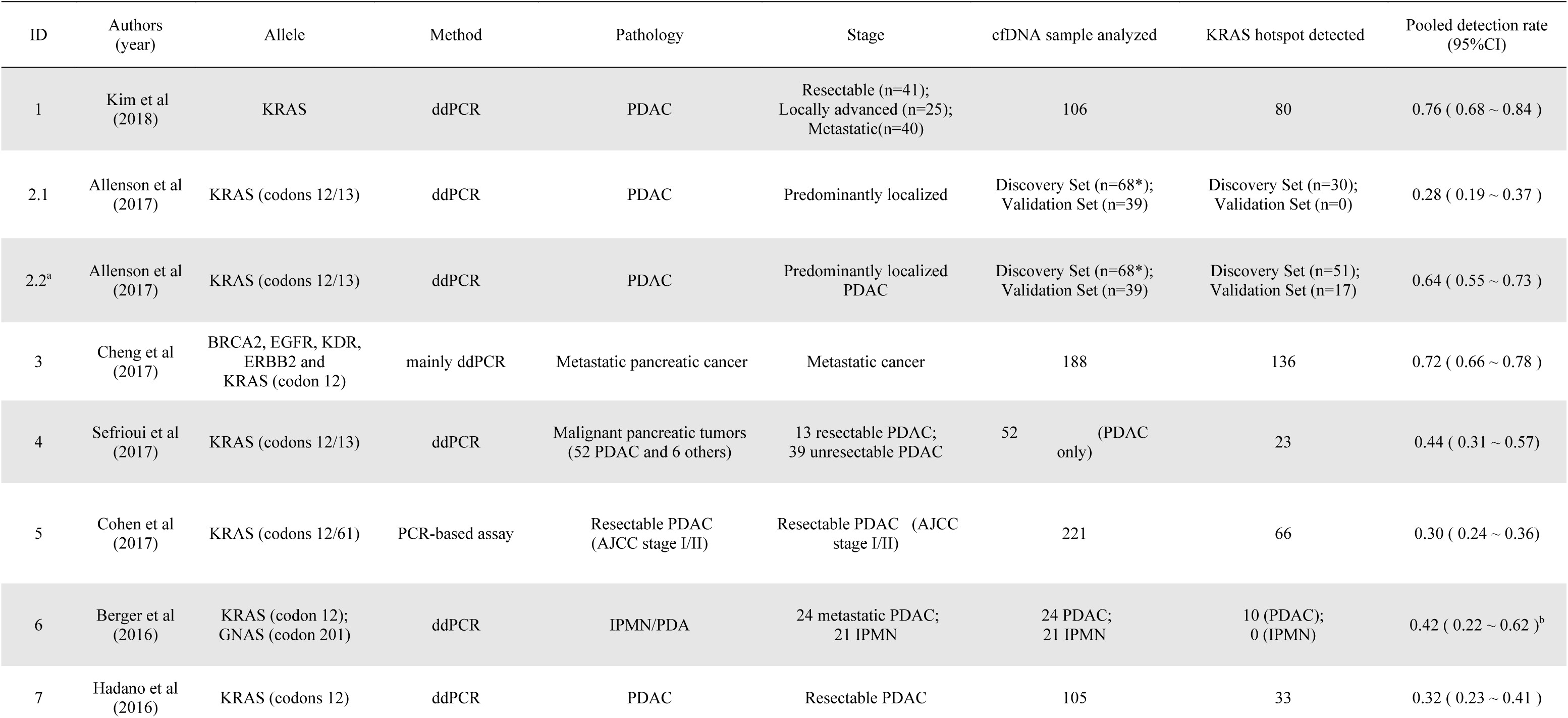
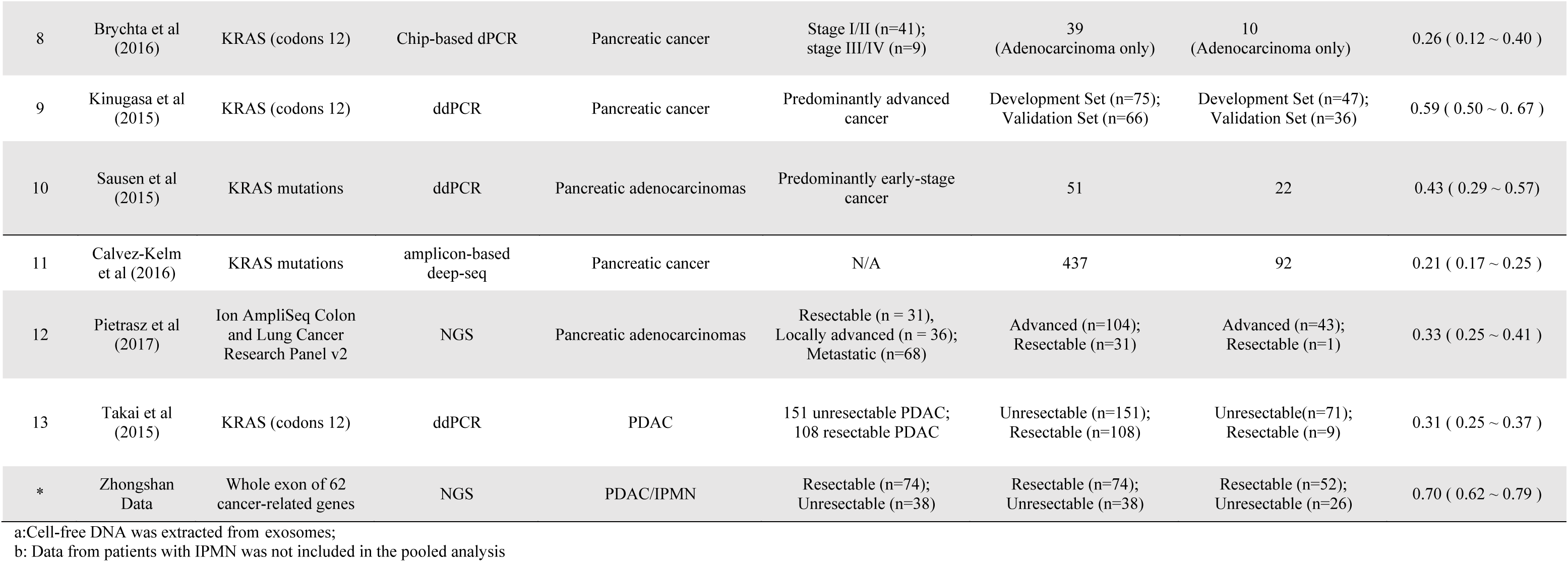
A meta-analysis of 13 high-quality studies measuring plasma KRAS mutations using PCR-or NGS-based methods.

**Table S5.** Univariate and multivariable survival analysis of the patient cohort.

| Clinical Characteristics | Univariate analysis |  | Multivariable analysis |  |
| --- | --- | --- | --- | --- |
|  | HR (95%CI) | *p*-value | HR (95%CI) | *p*-value |
| AJCC stage |  |  |  |  |
| Early stage (I/II) | 1.0 |  | 1.0 |  |
| Advanced (III/IV) | 8.4 ( 3.9 ~ 18.0 ) | <0.001*** | 3.7 ( 1.1 ~ 11.9 ) | 0.029* |
| CA19-9 (741.8 U/ml ) |  |  |  |  |
| Low | 1.0 |  | 1.0 |  |
| High | 4.6 ( 2.2 ~ 9.1 ) | <0.001*** | 1.3 ( 0.4 ~ 3.7 ) | 0.661 |
| CA125 (37.7 U/ml) |  |  |  |  |
| Low | 1.0 |  | 1.0 |  |
| High | 6.7 ( 2.9 ~ 15.6 ) | <0.001*** | 2.9 ( 0.9 ~ 10.2 ) | 0.087 |
| CEA (5.4 ug/ml) |  |  |  |  |
| Low | 1.0 |  | 1.0 |  |
| High | 3.13 ( 1.5 ~ 6.3 ) | 0.002** | 0.88 ( 0.3 ~ 3.1 ) | 0.843 |
| Total cfDNA (22.2 ng/ml)* |  |  |  |  |
| Low | 1.0 |  | NA |  |
| High | 3.57 ( 1.8 ~ 7.1 ) | <0.001*** | NA | NA |
| KRAS copies (86.4 copies/ul)* |  |  |  |  |
| Low | 1.0 |  | 1.0 |  |
| High | 4.35 ( 2.1 ~ 9.1 ) | <0.001*** | 3.3 ( 1.1 ~ 10.6 ) | 0.037* |
| KRAS fraction (0.68 %) |  |  |  |  |
| Low | 1.0 |  | NA |  |
| High | 2.12 ( 0.9 ~ 4.8 ) | 0.077 | NA | NA |

*These variables were not included in the multivariable survival analysis.

